# A new method to accurately identify single nucleotide variants using small FFPE breast samples

**DOI:** 10.1101/2020.10.22.350983

**Authors:** Angelo Fortunato, Diego Mallo, Shawn M. Rupp, Lorraine M. King, Timothy Hardman, Joseph Y. Lo, Allison Hall, Jeffrey R. Marks, E. Shelley Hwang, Carlo C. Maley

## Abstract

Most tissue collections of neoplasms are composed of formalin-fixed and paraffin-embedded (FFPE) excised tumor samples used for routine diagnostics. DNA sequencing is becoming increasingly important in cancer research and clinical management; however, it is difficult to accurately sequence DNA from FFPE samples. We developed and validated a new bioinformatic algorithm to robustly identify somatic single nucleotide variants (SNVs) from whole exome sequencing using small amounts of DNA extracted from archival FFPE samples of breast cancers. We optimized this strategy using 28 pairs of technical replicates. After optimization, the mean similarity between replicates increased 5-fold, reaching 88% (range 0-100%), with a mean of 21.4 SNVs (range 1-68) per sample, representing a markedly superior performance to existing algorithms. We found that the SNV-identification accuracy declined when there was less than 40ng of DNA available and that insertion-deletion variant calls are less reliable than single base substitutions. As the first application of the new algorithm, we compared samples of ductal carcinoma in situ (DCIS) of the breast to their adjacent invasive ductal carcinoma (IDC) samples. We observed an increased number of mutations (paired-samples sign test, p<0.05), and a higher genetic divergence in the invasive samples (paired-samples sign test, p<0.01). Our algorithm provides a significant improvement in detecting SNVs in FFPE samples over previous approaches.

**Key Points:**

- The sequencing of reduced quantities of DNA extracted from FFPE samples leads to substantial sequencing errors that require correction in order to obtain accurate detection of somatic mutations.
- We developed and validated a new bioinformatic algorithm to robustly identify somatic single nucleotide variants using small amounts of DNA extracted from archival FFPE samples of breast cancers.
- Variant calling software packages need to be optimized to reduce the impact of sequencing errors. Our bioinformatics pipeline represents a significant methodological advance compared to the currently available bioinformatic tools used for the analysis of small FFPE samples.

## Introduction

Tumors are characterized by a high genetic heterogeneity both within the same tumor type and in different parts of the same neoplasm [1]. Genetic heterogeneity determines the capacity of the neoplastic cell population to adapt to new microenvironments and to develop resistance to therapeutic treatments [2–4]. We and others have hypothesized that the quantification of genetic heterogeneity will be generally useful for risk stratification of patients [5,6]. However, in order to do so, we need accurate methods for identifying somatic genomic alterations in neoplasms. Cancers can develop from different combinations of genetic mutations and each patient typically has a unique mutational profile, distributed among a mosaic of subclones across the tumor [7]. This makes it difficult to develop universal biomarkers to predict cancer progression based on specific mutations and a single sample from a neoplasm. Alternatively, measures that characterize the underlying evolutionary process do not focus on specific progression mechanisms or the particular mutations that occur, making them more generalizable [6].

Intratumor heterogeneity is one such measure, and we have successfully used it in the past to predict cancer progression of pre-malignant diseases [8–10] and overall survival in cancers [3]. Routine diagnosis in oncology relies on histopathological analysis of formalin-fixed and paraffin-embedded (FFPE) excised tumor samples. Using these samples for genetic analysis has numerous advantages: histopathological analyses are already available for them, specific areas can be selected with precision eliminating the need to take additional samples dedicated to genetic analysis and, moreover, they are archived in large numbers, readily available to carry out retrospective studies. On the other hand, these samples have several technical limitations when used for genetic analyses. Histological fixation and embedding partially degrades and binds amino acids to the DNA, which continues to deteriorate over time [11]. Deamination of cytosine residues leading to apparent C to T transitions is also a common artefact in FFPE derived DNA [12]. These problems are exacerbated when the amount of available DNA is limited, because DNA artifacts are not compensated by the abundance of intact molecules, leading to sequencing errors [13,14]. This is particularly relevant when studying early or precancerous conditions where the lesion can be very small. In order to study genomic intratumor heterogeneity using FFPE samples, we must often sequence the degraded and imperfectly purified DNA extracted from small focal areas of the tumor or pre cancer. Furthermore, estimates of intratumor heterogeneity as well as other precision medicine efforts are confounded by both false positives and false negatives in the detection of mutations. Precision medicine requires avoiding false positives and negatives which would potentially expose patients to the wrong therapeutic interventions. Thus, there is a clear need for robust and accurate methods for sequencing and detecting mutations in small amounts of DNA extracted from FFPE samples. We have developed a new bioinformatic method that reduces these obstacles for the estimation of genetic intratumor heterogeneity using paired FFPE samples. We developed this somatic-variant post-processing pipeline by empirical optimization using 28 whole exome sequencing replicates—DNA samples sequenced twice independently, and validated the results using a different, high depth, sequencing technique.

Most scientific disciplines rely heavily on replication to measure stochasticity and reduce different types of errors. However, most sequencing experiments do not use any kind of biological or technical replication, relying on increasing levels of sequencing depth and post-processing strategies to improve their accuracy. This limitation has been highlighted in the past in a small number of studies [15,16]. These studies identified quality control metrics that correlate with the concordance between technical replicates and their relative importance.

However, only very recently has this concept been applied to the improvement of variant calling methods[17,18]. Karimnezhad et al. [17] advocate using the intersection SNVs identified by different methods and/or technical replicates, while Kim et al.[18] developed a variant calling method (RePlow) that leverages technical replicates to dramatically improve the specificity in the detection of somatic variants present at very low variant allele frequency. This approach is promising but requires the generation of technical replicates for all study samples, potentially doubling sequencing costs. Alternatively, here we present and implement a strategy to use a small number of technical replicates to optimize a pipeline, which then can be used to estimate intratumor genetic heterogeneity reliably without the need to use technical replicates for all study samples.

We selected a precursor of breast cancer, ductal carcinoma *in situ* (DCIS), to develop and optimize our pipeline because most of these tumors are detected in the early phase of their development, and there is an important clinical need to be able to estimate the risk level of this commonly diagnosed precancer in order to better understand the genomic changes that are associated with cancer progression. Improved risk stratification in DCIS could guide improvements in management of the condition and therapeutic intervention. The majority of breast tumors develop in the terminal duct lobular unit, mainly starting among duct cells [19,20] (Fig. 1). The cancer cells proliferate within the ducts and deform their anatomical structure. Despite the ducts’ growth in volume their walls remain intact, confining the tumor cells in the lumen, separating them from nearby tissues and limiting their dissemination. In this phase, the tumor is defined as ductal carcinoma *in situ* (DCIS). Subsequently, the cells may evolve to invasive disease, crossing the duct wall’s boundaries, invading the surrounding tissue, and potentially metastasizing. DCIS tumors can remain non-invasive but there is substantial evidence that a subset will invade and, in some cases, metastasize. The development of a new bioinformatic algorithm to identify somatic single nucleotide variants and measure genetic heterogeneity could provide a significant contribution to the estimation of DCIS patients’ risk for progressing to breast cancer.

**Figure 1:**
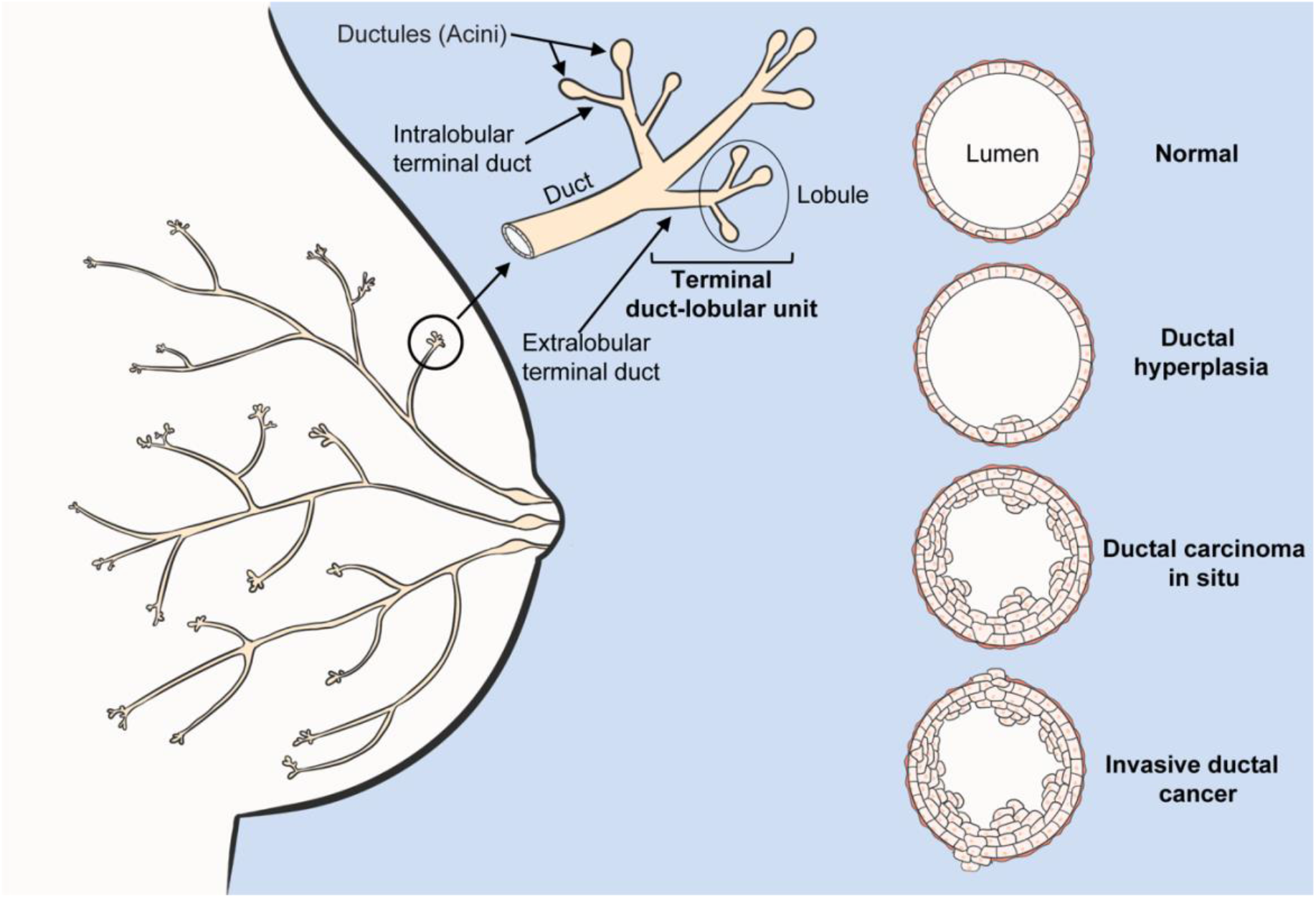
Breast cancer anatomy. Schematic representation of mammary gland anatomy and cancer development. The majority of breast tumors develop in the terminal duct lobular unit, 80% starting among ductal cells. Initially, the duct suffers a benign hypertrophic growth of cells that can progress into ductal carcinoma *in situ* (DCIS). In this phase the neoplasm is confined within the duct’s lumen and it is still clinically benign. Cancer cells can cross the duct wall’s boundaries, invading nearby tissues (IDC) and metastasizing.

## Results

Ideally, the same sample of tumor DNA, when sequenced twice with the same methodology, should give the same results (detect the same mutations). We developed our mutation detection pipeline (Fig. 2), optimized it using duplicate (technical replicate) whole exome sequencing of the same samples, and validated our results using deep targeted sequencing.

**Figure 2:**
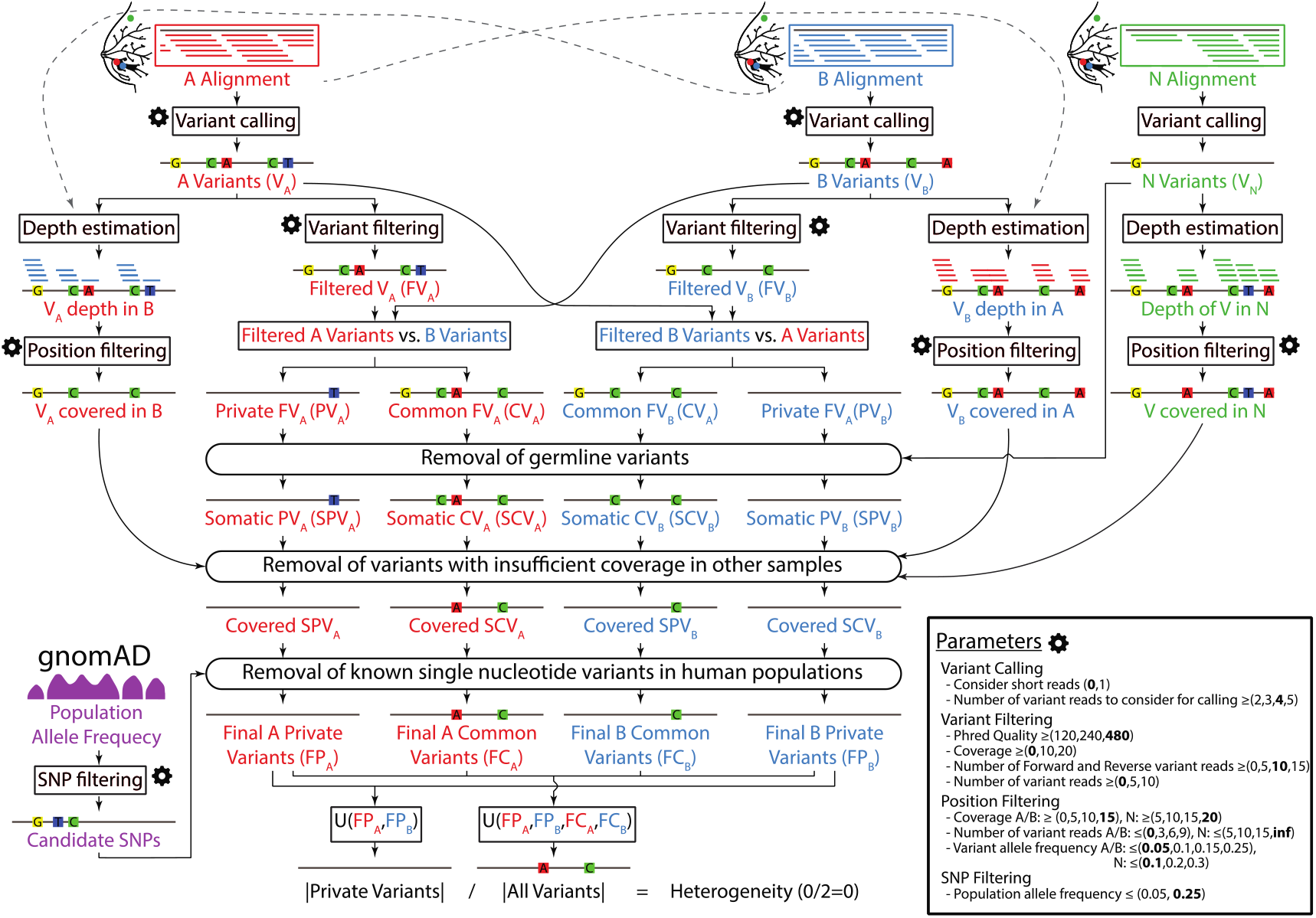
Flowchart of the algorithm used to estimate the genetic heterogeneity between two samples and details of its optimization. Inputs: aligned sequences (BAM files) of the two samples (A, in red; and B, in blue) and their healthy tissue control (N, in green), population allele frequency data from the gnomAD database (single nucleotide polymorphisms, SNPs, in purple), and user-specified configuration parameters (gear icon). Outputs: estimate of the genetic heterogeneity between samples A and B, and set of variants (level of detail user-specified). All parameters that control this pipeline are detailed in the Parameters box, accompanied by the range of values assayed during optimization between parentheses and the final set of optimized values in bold. The key

### Pipeline optimization

We used an empirical method for optimizing the analysis algorithm through the comparison of technical replicates of whole exome sequences. Any variant detected only in one sample but not in the other is likely the result of a sequencing or data processing error. This approach allowed us to systematically and objectively compare alternative parameterizations of the estimation pipeline to single out the best overall and to find the most generalizable parameter values using cross-validation.

In order to optimize our pipeline, we assigned a range of values to explore for each of the 13 parameters that control its execution (Fig. 2) and explored every possible combination of them, scoring each using a statistic that integrates the central tendency and dispersion of the heterogeneity across the 28 technical replicates. Furthermore, we used DNA quantity (from 20 ng to >100 ng) in order to evaluate the efficiency of the method on different quantities of input DNA, in order to determine the limits of the method on small amounts of DNA (Suppl. table 1S). The resulting algorithm (Fig. 2) yielded a mean similarity across the 28 technical replicates of 88% (range 0-100%) (Fig. 3), which constitutes a 5-fold improvement over using the same variant caller–Platypus without any post-processing of the results [21], (17.8%, range: 0.1-61.8%). We identified a mean of 21.4 (range 1-68) single nucleotide variants per sample (Table 1), which are distributed throughout the entire exome (Fig. 3S).

**Table 1:** Similarity between technical replicates and number of variants. The similarity between technical replicates on average is 88%, range 0-100%. Number of total, common and private SNVs (A+B). Common SNVs: SNVs detected in both replicas of the same DNA samples; Private SNVs: SNVs detected only in one of the two DNA sequences of the same DNA.

| Sample | Common | A + B | Total | Similarity (%) |
| --- | --- | --- | --- | --- |
| DCIS-017 | 0 | 1 | 1 | 0 |
| DCIS-020-B3 | 19 | 8 | 27 | 70.4 |
| DCIS-020-B6 | 57 | 11 | 68 | 83.8 |
| DCIS-028-K12 | 4 | 0 | 4 | 100 |
| DCIS-029-D5 | 20 | 6 | 26 | 76.9 |
| DCIS-029-D8 | 11 | 2 | 13 | 84.6 |
| DCIS-050 | 8 | 1 | 9 | 88.9 |
| DCIS-064 | 28 | 2 | 30 | 93.3 |
| DCIS-080 | 7 | 0 | 7 | 100 |
| DCIS-094-B11 | 45 | 4 | 49 | 91.8 |
| **DCIS-094-B7** | 35 | 1 | 36 | 97.2 |
| **DCIS-122** | 3 | 0 | 3 | 100 |
| **DCIS-135** | 9 | 2 | 11 | 81.8 |
| **DCIS-163** | 1 | 0 | 1 | 100 |
| **DCIS-164** | 44 | 2 | 46 | 95.7 |
| **DCIS-168-C4** | 55 | 2 | 57 | 96.5 |
| **DCIS-168-C8** | 41 | 0 | 41 | 100 |
| **DCIS-171** | NA | NA | NA | NA |
| **DCIS-178** | 8.0 | 0 | 8 | 100 |
| **DCIS-211** | 12 | 0 | 12 | 100 |
| **DCIS-213** | NA | NA | NA | NA |
| **DCIS-222-B10** | 6 | 0 | 6 | 100 |
| **DCIS-222-B6** | 1.0 | 0 | 1 | 100 |
| **DCIS-225-A16** | 9 | 5 | 14 | 64.3 |
| **DCIS-225-A6** | NA | NA | NA | NA |
| **DCIS-227** | 6 | 0 | 6 | 100 |
| **DCIS-250** | NA | NA | NA | NA |
| **DCIS-267** | 33 | 5 | 38 | 86.8 |
| **Average** | **19.3** | **2.2** | **21.4** | **88.0** |
| **S.D.** | **18.2** | **2.9** | **19.8** | **21.4** |

**Figure 3:**
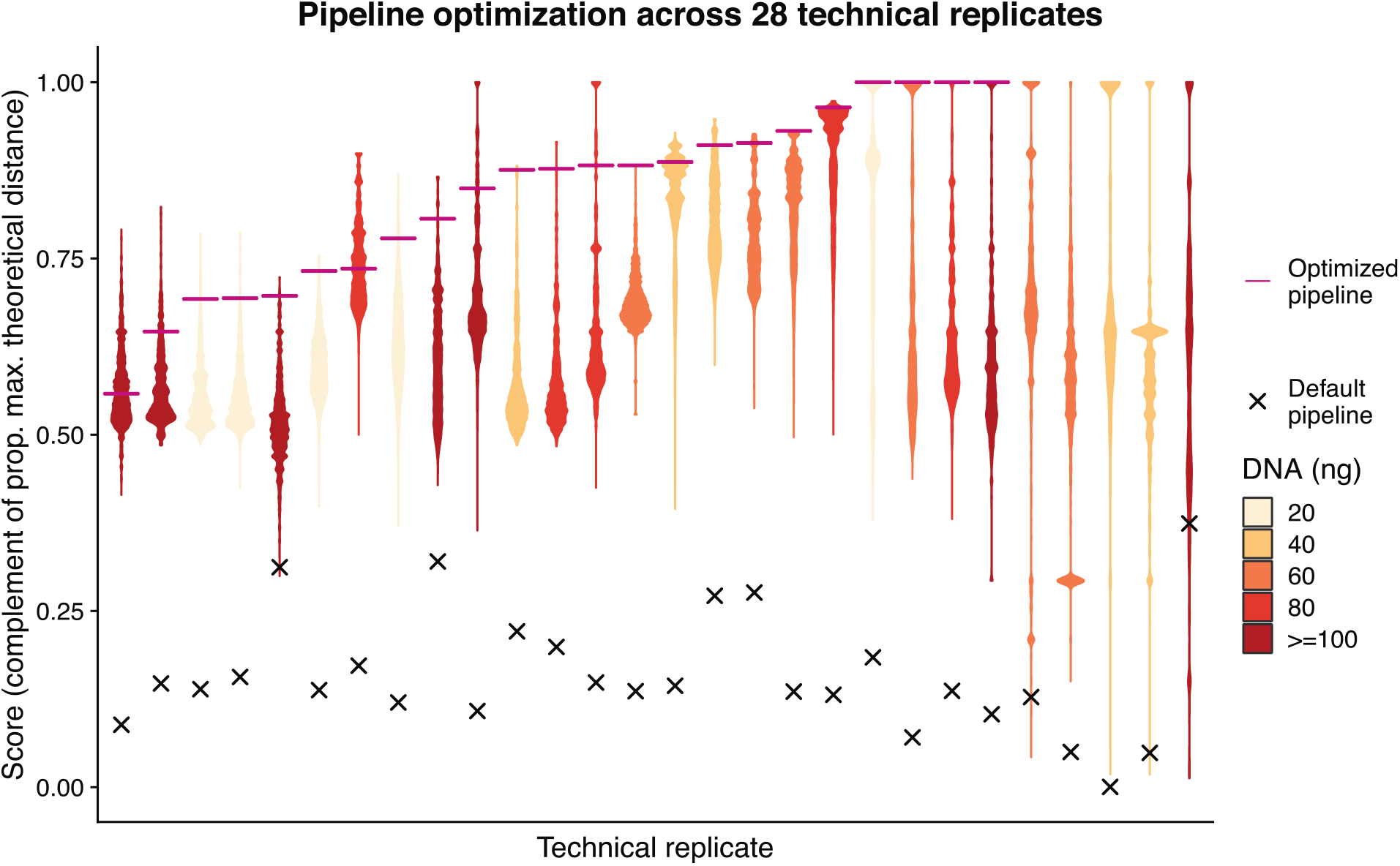
Empirical optimization of the variant post-processing algorithm. Each violin plot summarizes the distribution of optimization scores of 5,308,416 combinations of values of the 13 parameters that control the pipeline for one of the 28 technical replicates (same DNA sample processed twice independently). The optimization score indicates the two-dimensional euclidean distance to the theoretical optimum value of similarity between technical replicates (1) and proportion of final common variants that have a population allele frequency below 0.05 (1) relative to the maximum possible distance. After parameter optimization the similarity between the technical replicates was on average 88 %, range 0-100% (x= score before optimization; **—**: score after optimization; colors indicate the amount (ng) of DNA used as template).

We also assayed an alternative implementation of our algorithm that uses Mutect2 to call variants, but it achieved a much lower accuracy, with a mean similarity (including indels) across the 28 technical replicates of only 2.4%, range 0.4-6.9%. Overall, we found that only 14.9% of the single nucleotide variants overlap between our main pipeline and this alternative implementation using Mutect2.

### Intratumor genetic heterogeneity estimation pipeline

In order to estimate the genetic heterogeneity between two samples (A, B), we applied the concept that the presence of a high confidence variant in one sample should increase the confidence of that variant in the other sample. This concept could also be applied to multi-region sequencing projects. We implemented this in a crossed unequal comparison scheme (Fig. 2), by which the set of filtered variants detected in a sample is compared against all variants estimated in the other sample. This comparison is then reversed, to finally integrate the result of the two comparisons by considering any variant found common in either comparison as common, or private otherwise. Thus, if a variant has been detected with high confidence in one sample and has also been detected in the other sample–even if with low confidence–the variant is considered present in both samples. However, if a variant is detected with low confidence in both samples the variant is discarded, preventing an artificial increase in the confidence of shared variants. Finally, variants that are detected with high confidence in only one sample and not detected even at low confidence in the other sample, are considered private. Before the integration step, the algorithm refines the variants removing detected germline variants, known germline variants in human populations, and variants with insufficient coverage in either the normal sample (all variants) or the other sample (private variants) (see Methods for additional details).

### Validation of filtering parameters

We performed a 5-fold cross-validation study to assess the sensitivity of the optimization strategy to input data, and how well the algorithm generalizes to independent datasets. The optimization strategy was relatively robust to the input data, returning a mean evaluation score (empirical cumulative distribution of test score) of 0.79, range 0.4 - 1 (Suppl. fig. 1S). Importantly, this experiment shows the robustness of the overall optimal model across different cross-validation folds, being the model with the highest mean training score and within the top 0.00006% of the mean test scores in this cross-validation analysis. The test score of the overall optimal model is always as good or better than the model selected based on the training score for each fold.

### Sensitivity analysis of the number of technical replicates

We saw a fast increase in the relative score, reaching a plateau with just 6 technical replicates and exhibiting diminishing returns when going over 10 technical replicates (Suppl. fig. 2S). With 6 technical replicates the results are very close to the ones obtained using the whole dataset, resulting in conditions that show a mean empirical cumulative probability of the optimization score that is 0.98 times the score obtained using all samples.

### Validation of somatic variants

In order to validate the identified mutations with our new method, we analyzed the same DNA used for the exome sequences using targeted primers and the AmpliSeq™ technology. We achieved an average of 18,821 (tumor) and 12,904 (control) read coverage for each single nucleotide variant in the validation set. The comparison of the data confirmed 89.6% (with optimal parameters, O) and 86.3% (with permissive parameters, P) single nucleotide variants identified by applying our pipeline to the exome sequence (Table 2). We found 2 (O) or 2 (P) of the unconfirmed variants belong to the same gene MUC6 characterized by highly repetitive sequences, thus subject to read alignment errors and known to have an unreliable reference sequence [22]. Excluding all MUC6 (3 (O) or 3 (P) variants), we validated 90.7% (O) or 86.7% (P) of the remaining variants. We found that 21.4% (O) and 18.7 % (P) of the confirmed variants are also present in the control samples with a frequency >10%; thus, these could be SNPs and not somatic mutations (Table 2). However, the expected frequency (50%) of the two alternative alleles of a germline SNP only occurs in 7 (O and P) cases, if we include alleles with frequency >40% (Suppl. table 2S, Fig. 4S). Importantly, we found a strong negative correlation between the amount of input DNA used (20, 40, 60 and 80 ng, validation set) for the NGS libraries and the inability to identify correctly the SNPs in the germ line DNA (Spearman correlation r = −0.31, p<0.0001(O), r=-0.28, p<0.001(P); Suppl. table 2S). Excluding MUC6 variants and DNA samples with less than 40 ng, we validated 94.7% (O) or 93.2% (P) of the variants, however, 3 (2.7%) (O) or 3 (2.3%) (P) variants were detected only in one of the two technical replicates.

**Table 2:** Validation. Targeted sequencing confirmed that 89.6% (Optimal filtering pipeline) and 86.3% (Permissive filtering pipeline) of single nucleotide variants identified using our algorithm. Excluding MUC6 and low input amounts of DNA we validated 94.7% (O) or 93.2% (P) of variants. We found that the 14.2% (O) or 12% (P) of the confirmed variants are also present in the control samples with a frequency >10%. These variants may be SNPs.

| **Optimal filter (O)** | **Variants** | **Common (A and B)** | **Private (A or B)** | **Variants in controls (>10%)** |
| --- | --- | --- | --- | --- |
| **Total number of SNVs** | 154 | 146 (94.8%) | 8 (5.2%) | 33 (21.4%) |
| Validated variants | 138 (89.6%) | 133 (91.1%) | 5 (62.5%) | 32 (97%) |
| Non-validated variants | 16 (10.4%) | 13 (8.9%) | 3 (37.5%) | 1 (3%) |
| **MUC6-excluded, DNA ≥ 40ng SNVs** | 113 | 110 (97.3%) | 3 (2.7%) | 16 (14.2%) |
| Validated variants | 107 (94.7%) | 105 (95.5%) | 2 (66.7%) | 15 (93.8%) |
| Non-validated variants | 6 (5.3%) | 5 (4.5%) | 1 (33.3%) | 1 (6.3%) |
| **Permissive filter (P)** | **Variants** | **Common (A and B)** | **Private (A or B)** | **Variants in controls (>10%)** |
| **Total number of SNVs** | 182 | 170 (93.4%) | 12 (6.6%) | 34 (18.7%) |
| Validated variants | 157 (86.3%) | 152 (89.4%) | 5 (41.7%) | 33 (97.1%) |
| Non-validated variants | 25 (13.7%) | 18 (10.6%) | 7 (10.6%) | 1 (2.9%) |
| **MUC6-excluded, DNA ≥ 40ng SNVs** | 133 | 130 (97.7%) | 3 (2.3%) | 16 (12%) |
| Validated variants | 124 (93.2%) | 122 (93.8%) | 2 (66.7%) | 15 (93.8%) |
| Non-validated variants | 9 (6.8%) | 8 (6.2%) | 1 (33.3%) | 1 (6.3%) |

**Figure 4.**
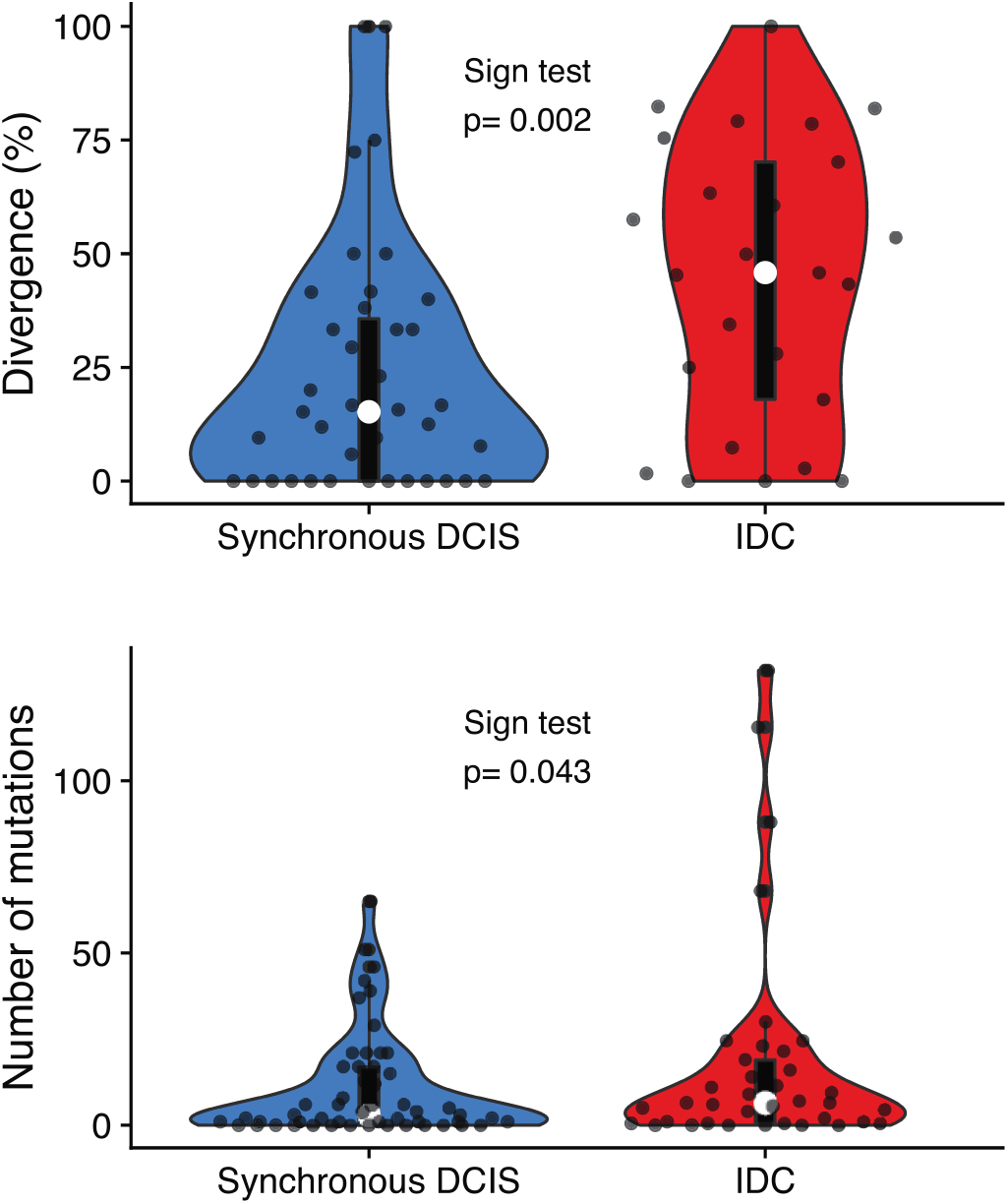
Mutational burden and genetic divergence. The average of the number of mutations of synchronous DCIS samples (10.40±15.31 S.D.) is lower than the IDC samples (18.05±31.48 S.D.) and there is a statistically significant difference between the two groups, paired-samples sign test, p<0.05. We found a statistically significant difference in genetic divergence comparing two regions of synchronous DCIS (21.48%±17.54 S.D.) versus the divergence between synchronous DCIS IDC samples (44.51%±29.04 S.D.) within the same patient, paired-sample sign test and Mann-Whitney U test, p<0.01. White circle=median, box limits indicate the 25th and 75th percentiles; whiskers extend 1.5 times the interquartile range from the 25th and 75th percentiles; curves represent density and extend to extreme values. Data points are plotted as dots.

We found that insertion-deletion variants are an unreliable sub-set of mutations (22 (O) and 16 (P) indels tested: 31.8% (O) and 31.3 (P) indels fully validated, 31.8% (O) and 25 (P) indels partially validated, in which not all nucleotides have been confirmed).

### Breast cancer genetic divergence

In order to showcase the application of our algorithm, we compared synchronous samples from two regions of DCIS and one sample of invasive ductal carcinoma (IDC) in each of 53 patients. We found a statistically significant difference in the number of mutations between these two diseases, (mean 10.40 in DCIS and 18.05 in IDC, paired-samples sign test, p<0.05). Importantly, our method allowed us to measure a statistically significant genetic divergence (heterogeneity) between the two synchronous DCIS samples and between DCIS vs. IDC samples (Fig. 4) (paired-samples sign test, p<0.01; Mann-Whitney U test, p<0.01). Genetic divergence is defined as the percentage of mutations detected in the union of the mutations from the two samples that are not shared by both samples. It is a common metric in evolutionary biology to estimate the amount of evolutionary change that has occurred since two populations shared a common ancestor. Previous work has shown that genetic divergence can predict progression to malignancy [8–10].

## Discussion

Cancer is a disease of clonal evolution, and intra-tumor heterogeneity is its fuel. There is increasing recognition that this heterogeneity poses a challenge for traditional sampling and prognosis, as different biopsies may sample different clones with variable relevance to the future behavior of the tumor. However, because heterogeneity itself drives clonal evolution, the magnitude of heterogeneity may itself be prognostic. Our previous studies of metrics of intratumor heterogeneity, showed that one robust measure is the degree to which two samples from the same tumor have genetically diverged (i.e. genetic diversity) [9]. This measure has the useful property that the more of the genome that is sequenced, the more accurate it becomes. We hypothesized that those ductal carcinoma *in situ* (DCIS) lesions with greater clonal heterogeneity would be more likely to progress to invasive and metastatic disease. However, in order to test that hypothesis, we required a reliable method to measure clonal heterogeneity in this experimental system. Here we have developed, characterized, and validated a method to measure genetic divergence from two FFPE derived DNA samples from the same tumor, solving this limitation. Our bioinformatics pipeline represents a significant methodological advance compared to the currently available bioinformatic tools used for the analysis of small FFPE samples.

The sequencing of small quantities (less than 200ng) of DNA extracted from FFPE samples leads to low coverage, high duplication rates, and substantial sequencing errors that require correction in order to obtain accurate detection of somatic mutations. Variant calling software packages need to be optimized to reduce the impact of sequencing errors. This is particularly important in the study of heterogeneity, as well as precision medicine, as both false positive and false negative detection of mutations can impact clinical decision making and diminish the predictive power of heterogeneity as a potential biomarker.

Any study of tumor heterogeneity using comparable DNA samples must account for and minimize technical variation. We found 88% of the variants were detected in both duplicated sequences and 94.7% excluding the MUC6 gene and those samples with ≤40ng input DNA. Both levels of filtering stringency tested (Optimal and Permissive) have proven successful. As expected, the relaxed version of the algorithm allows the detection of a higher number of variants in exchange for a small reduction of accuracy. It is surprising that, when not using a post-processing pipeline such as the one presented here, variant callers like Platypus and Mutect2 generated very inaccurate results on our WES data, with similarities between the technical replicates of only 17.8% and 2.4%, respectively. Our systematic study reveals the magnitude of uncertainty related to making mutation calls from small amounts of FFPE derived DNA.

We validated the bioinformatic algorithm by re-sequencing the regions containing the variants using a different sequencing technique: AmpliSeq™. This technology allows for a deep re-sequencing of the regions of interest, improving our ability to identify mutations correctly. The comparison between the data obtained with these two techniques allowed us to validate the new algorithm. Among these, some are presumably SNPs and not somatic variants. However, the frequency of the two alternative alleles is often far from the expected frequency of 50%. This could be because of difficulties encountered when sequencing with AmpliSeq™ to analyze DNA extracted from FFPE, or biological signals of neoplastic DNA present in the control samples.

The fact that there is a strong statistically significant negative correlation between the amount of DNA used for the preparation of the libraries and the presence of SNPs detected as SNVs suggests that at least 40 ng of input DNA be used for standard library preparation. In particular, this result indicates that the quality and quantity of control DNA is a key factor in the ability to correctly identify somatic mutations in tumors. In many instances, control DNA is not a limiting factor and higher amounts can be used for the preparation of the NGS libraries. Moreover, control samples could be collected during surgery or from blood cells, obtaining DNA from specimens that have not undergone the effect of fixation and DNA deterioration. Our algorithm allows us to modulate the stringency of SNP filtering parameters and to obtain the frequency of each potential SNP in the population.

The variants detected using our algorithm were distributed over the entire exome and we have cataloged numerous mutations in well-known breast cancer genes. As a first application of the new algorithm, we compared synchronous DCIS and invasive (IDC) samples. We identified a statistically significant increase in the number of mutations and genetic divergence in the invasive samples compared to DCIS samples. This result has been described in other types of tumors [9]. Given these findings, we can test if genetic divergence between regions of DCIS predicts future recurrence of DCIS or progression to IDC in a larger cohort.

The current version of our algorithm has been developed and implemented to fit our needs, analyzing two samples per patient to measure their genetic divergence. However, this strategy is easy to generalize to any number of samples to apply it to larger multi-region datasets. We have not done it here since there are some nuances that may need to be adjusted depending on the final purpose of the called SNVs. The removal of variants with insufficient coverage in other samples is the main focus of these decisions. For example, for a downstream analysis that does not integrate uncertainty easily, the algorithm could require enough coverage in most (or all) samples, discarding variants with a lot of missing data, while for other applications those SNVs could be kept if they are at least present in another sample, assigning missing values or a measure of uncertainty to samples with insufficient coverage. The core step of the algorithm— comparison of filtered and unfiltered sets of variants—could be kept as it is. However, we also envision more stringent alternatives in which a variant must be present in more than one non-filtered sample to be kept in the final set. The removal of germline variants and SNPs would remain, since it does not depend on the number of samples.

## Conclusion

We developed a bioinformatics pipeline to analyze pairs of DCIS samples taken from the same neoplasm. We identified the mutations present in each sample and we showed that this method has high fidelity in technical replicates and is capable of identifying different levels of genetic heterogeneity between regions of the same tumor. This algorithm is easily modifiable and can be integrated with additional parameters, allowing investigators to choose different levels of filtering stringency. These parameter values can be re-optimized for a different experimental system with as few as six sets of technical replicates, and the optimized set of parameter values provided here is robust to changes in the input data and thus is expected to translate well to other systems. These characteristics make our algorithm readily applicable to large tissue banks of FFPE samples of any neoplasm and is particularly useful for studies to quantify genomic heterogeneity.

## Methods

### Patients clinical data and biological samples

This study was approved by the Institutional Review Board (IRB) of Duke University Medical Center, and a waiver of consent was obtained according to the approved protocol. Formalin-fixed paraffin embedded (FFPE) breast tissue blocks were retrieved from Duke Pathology archives. All cases underwent pathology review (AH) for tissue diagnosis and case eligibility.

Breast tumors were classified using the World Health Organization (WHO) criteria [23]. Following pathology review, a total of 66 separate patients are included in this study. All DNA was extracted from archival formalin fixed paraffin embedded thin sections stained with hematoxylin. For tumors, the study pathologist identified areas of DCIS or invasive cancer that were macrodissected to enrich for tumor epithelial cells. Control DNA was extracted from either distant benign areas of the breast or a benign lymph node using the same procedure employed for the tumor containing areas. These benign areas were confirmed to be devoid of tumor by the study pathologist.

A total of 28 breast tumor DNA samples were included in the development of the method procedure divides as follows: pure DCIS (DCIS not associated with invasion; n=15 tumors, from 11 patients), synchronous DCIS (DCIS identified concurrently with invasive cancer; n=6 tumors, from 6 patients) and invasive ductal carcinoma (IDC; n=7 tumors, from 5 patients) (Table 3). 53 synchronous DCIS patients were used for the experimental validation of the new algorithm. For each patient we selected two DCIS samples located at least 8mm apart (total 106 samples) and 37 IDC samples derived from the same synchronous DCIS patients. Each specimen was macrodissected and DNA extracted separately. IDCs and DCIS were graded according to the Nottingham grading system [24] or recommendations from the Consensus conference on DCIS classification [25], respectively.

**Table 3:** Patients clinical data. Clinical data of the 22 patients included in the study. The histopathological analysis showed that 11 patients are DCIS while 6 are DCIS adjacent to invasive disease (DCIS Adj. to IDC) and 5 have invasive features (IDC). We selected FFPE samples of different ages (1999-2017). ER: estrogen receptors, PR: progesterone receptors, HER2: human epidermal growth factor receptor 2 expression is qualitatively estimated (non-present (NP), 0-8) using histochemistry stains.

| Patient ID | Age | Race | Date | Tumor type | Histopathological classification | ER | PR | HER2 | DCIS Size (mm) | DCIS nuclear grade | Invasive present |
| --- | --- | --- | --- | --- | --- | --- | --- | --- | --- | --- | --- |
| DCIS-017 | 66 | B | 2013 | Pure DCIS | cribriform, solid | - | + | NA | 21 | 3 | No |
| DCIS-020-B3 | 67 | W | 2014 | Pure DCIS | cribriform, solid, micropapillary, comedo | + | + | NA | 40 | 2 | No |
| DCIS-020-B6 | 67 | W | 2014 | Pure DCIS | cribriform, solid, micropapillary, comedo | + | + | NA | 40 | 2 | No |
| DCIS-029-D5 | 34 | W | 2012 | Pure DCIS | comedo | + | + | NA | 83 | 3 | No |
| DCIS-029-D8 | 34 | W | 2012 | Pure DCIS | comedo | + | + | NA | 83 | 3 | No |
| DCIS-050 | 52 | W | 2010 | Synchronous DCIS | cribriform, solid | + | + | - | 10 | 2 | Yes |
| DCIS-064 | 50 | OTHER | 2015 | Synchronous DCIS | comedo | + | + | + | 75 | 3 | No |
| DCIS-080 | 49 | W | 2013 | Synchronous DCIS | solid, comedo | + | + | - | 21 | 3 | Yes |
| DCIS-094-B11 | 68 | W | 2013 | IDC | cribriform, solid, micropapillary | - | - | - | NA | 3 | Yes |
| DCIS-094-B7 | 68 | W | 2013 | IDC | cribriform | - | - | - | NA | 3 | Yes |
| DCIS-122 | 47 | W | 2002 | Pure DCIS | cribriform, solid, comedo | NA | NA | NA | 95 | 3 | No |
| DCIS-135 | 48 | B | 2013 | Pure DCIS | cribriform, solid | + | + | NA | 13 | 2 | No |
| DCIS-163 | 53 | W | 2013 | Synchronous DCIS | cribriform, solid, comedo | + | + | - | 54 | 3 | Yes |
| DCIS-164 | 65 | B | 2015 | IDC | micropapillary, comedo | + | + | - | NA | 3 | Yes |
| DCIS-168-C4 | 63 | W | 2016 | IDC | cribriform, solid | + | + | - | NA | 2 | Yes |
| DCIS-168-C8 | 63 | W | 2016 | IDC | cribriform, solid | + | + | - | NA | 2 | Yes |
| DCIS-171 | 66 | B | 2000 | Synchronous DCIS | solid | - | + | - | 15 | 3 | Yes |
| DCIS-178 | 56 | W | 2011 | Synchronous DCIS | comedo, solid, micropapillary, papillary | - | - | - | NA | 3 | Yes |
| DCIS-211 | 43 | H | 2011 | Pure DCIS | cribriform, solid, comedo | + | + | NA | 24 | 3 | No |
| DCIS-213 | 68 | W | 2009 | Pure DCIS | cribriform, micropapillary, comedo | + | + | NA | 16 | 3 | No |
| DCIS-222-B10 | 41 | A | 2013 | Pure DCIS | cribriform, papillary | + | + | NA | 40 | 2 | No |
| DCIS-222-B6 | 41 | A | 2013 | Pure DCIS | cribriform, papillary | + | + | NA | 40 | 2 | No |
| DCIS-225-A16 | 62 | B | 2011 | Pure DCIS | cribriform, solid | + | + | NA | 30 | 2 | No |
| DCIS-225-A6 | 62 | B | 2011 | Pure DCIS | cribriform, solid | + | + | NA | 30 | 3 | No |
| DCIS-227 | 75 | B | 2012 | Pure DCIS | cribriform, solid, comedo | + | + | NA | 63 | 3 | No |
| DCIS-250 | 56 | W | 1999 | IDC | cribriform, comedo | + | - | NA | NA | 3 | Yes |
| DCIS-267 | 66 | W | 2017 | IDC | solid | + | + | - | 13 | 3 | Yes |
| DCIS-28-K12 | 42 | A | 2014 | Pure DCIS | comedo, micropapillary | - | - | NA | 124 | 3 | No |

### DNA extraction

The DCIS component of all cases as well as IDC from synchronous DCIS cases were macrodissected separately, following hematoxylin staining, of between 10 and 25 five-micron-thick histological sections. The first and last slides were stained with hematoxylin-eosin (H&E) staining and reviewed by a pathologist to confirm the presence of >=70% of neoplastic cells.

DNA was extracted using the FFPE GeneRead DNA Kit which incorporates enzymatic cleavage of DNA at uracil residues via uracil DNA glycosylase reducing the problem of cytosine deamination (Qiagen, cat n. 180134) according to manufacturers’ instructions. DNA quantification was performed using a Qubit™ 1X dsDNA HS Assay Kits (ThermoFisher, cat. n. Q33230), and DNA quality assessed with an Agilent 2100 Bioanalyzer.

### DNA sequencing

We sequenced different quantities of genomic DNA (20, 40, 60, 80, 100, >100 ng) to estimate the effects of DNA quantity on the estimation of intratumor genomic heterogeneity. All technical replicates were separated into two aliquots from the same tube of DNA sample before all subsequent steps. For experimental validation of the new algorithm, we used ≥40 ng of genomic DNA. Each aliquot was sheared to a mean fragment length of 250 bp using the Covaris LE200 instrument, and Illumina sequencing libraries were generated as dual-indexed, with unique barcode identifiers, using the Accel-NGS 2S PCR-Free library kit (Swift Biosciences, cat. n. 20096). We pooled groups of 96 equimolar libraries (100 ng/library) for hybrid capture using two target panels, the human exome and a panel containing all exons of the 83 genes in the breast cancer gene panel (BRC83, Suppl. table 4S). To capture BRC83 we used biotinylated “ultramer” oligonucleotides synthesized by Integrated DNA Technologies (Coralville, Iowa), and to capture the human exome we used IDT’s xGen Exome Research Panel v1.0. After hybridization, capture pools were quantitated via qPCR (KAPA Biosystems kit). We sequenced the final product using an Illumina HiSeq 2500 1T instrument multiplexing nine tumor samples per lane.

After binning the sample data according to its index identifier, we aligned it to the Genome Reference Consortium Human Build 37 using the BWA-MEM (Li, 2013) algorithm, and marked sequencing duplicates with Picard’s MarkDuplicates. The resulting BAM files are the input data for our pipeline for intratumor genetic heterogeneity calculation. We discarded samples with less than 40% of the target covered at 40X (Suppl. table 1S). This sequencing protocol was performed at the McDonnell Genome Institute at Washington University School of Medicine in St. Louis.

### Intratumor genetic heterogeneity estimation pipeline

We implemented our heterogeneity estimation pipeline (Fig. 2) in a series of Perl scripts, tailored to be run at Arizona State University’s research computing high performance computing clusters. Variants are first called using Platypus 0.8.1 [21] against the Genome Reference Consortium Human Build 37 reference genome using the default settings except for the parameters regulated during pipeline optimization (Fig. 2): The inclusion of reads with small inserts (-- filterReadPairsWithSmallInserts), and the minimum number of reads supporting a variant to consider it for calling (--minReads). Before downstream analyses, our pipeline splits multiallelic sites into biallelic sites, and clusters of variants into individual SNVs. The variant filtering step uses SnpSift 4.2 [26] (Phred Quality: QUAL, Coverage: GEN[*].NR[*], Forward and Reverse variant reads: NF & NR, Variant reads: GEN[*].NV[*]). The depth estimation step, which estimates the coverage of the position of a variant in the other samples (and the proportion of reads supporting that specific allele) is carried out by first generating a bed file integrating deletions, insertions, and SNVs using BEDOPS [27], and then using it as intervals input for GATK 3.5.0’s UnifiedGenotyper, executed to output data for all sites (--output_mode EMIT_ALL_SITES, -glm BOTH). The position filtering step is carried out in the inhouse pipeline with these results. This step differs slightly in the comparison between tumor samples and the comparison against the normal. In the first case, a variant is discarded if any of the conditions is not met, while in the second both the allele frequency and the number of variants need not be met for them to trigger the discard of a variant while the coverage filter acts independently. Importantly, while the steps of variant removal are generally applied to all sets of variants (e.g., removal of germline variants, candidate SNPs, and positions with lack of support in the normal), the removal of variants based on insufficient coverage in the other tumor samples only applies to private variants.

Population allele frequency estimates are obtained from the gnomAD 2.1.1 genomic database [28], which spans 15,708 whole-genome sequences, and filtering using this information is carried out within our pipeline. All variant comparisons within our pipeline are genotype specific.

We also implemented an alternative version of this pipeline identifying somatic mutations using Mutect2 [29] version 4.0.5.0 for comparison purposes against a developing version of our pipeline, both lacking the population allele frequency step (Fig. 2), and using slightly different parameter values, which were optimal at that stage of development (Suppl. table 5S). To use this variant caller, first we generated a panel of normals using all control tissue samples and the CreateSomaticPanelOfNormals GATK command. Then, we called variants on all paired tumor files using the panel of normals, IDT’s xGen Exome Research Panel v1.0, and the AllowAllReadsReadFilter. We filtered the resulting variants with an equivalent re-implementation of our post-processing pipeline that uses Bcftools isec to perform comparisons between sets of variants and ran FilterMutectCalls to obtain the final calls.

### Optimization of the intratumor genetic heterogeneity pipeline

We assigned a range of values to explore for each of the 13 parameters that control the genetic heterogeneity estimation pipeline (Fig. 2) and explored every possible combination of them with the data from all 28 technical replicates, assessing a total of 5,308,416 parameter combinations. We calculated the score of a condition (set of parameter values) as the minimum value of the 90% confidence interval of the mean (p=0.9) of the scores of that condition across the 28 technical replicates. We used this statistic to integrate central tendency and dispersion in the same measure. The score of each technical replicate was calculated as the two-dimensional euclidean distance to the theoretical optimum value of similarity between technical replicates (1) and proportion of final common variants that have a population allele frequency below 0.05 (1) relative to the maximum possible distance. This score ranging from 0 to 1, allowed us to co-optimize the similarity between technical replicates and the sets of variants with the least chance of being dominated by germline variants not detected in the normal and detected as somatic common variants. We performed a 5-fold cross-validation study stratified by amount of DNA, in which patients were partitioned randomly into 5 subsets, with at least 1 patient from each DNA category 20, 40, 60, 80, ≥100 ng. In each of the 5 interactions, one of the subsets (testing set) was held out of the parameter optimization and then evaluated based on the optimal parameter values obtained from the training set. We implemented the optimization and cross-validation steps in R [30], using the LSR (Navarro 2015), and cowplot [31] packages.

### Sensitivity analysis on the number of technical replicates

We subsampled our dataset to create smaller technical replicate datasets of k={2,…,28} sizes. For each k, we generated all combinations of size k with our 28 technical replicates and took a random sample of 104 of them (or all if ≤104) without replacement. We optimized the pipeline using each of these resampled subsets and reported the empirical cumulative probability of its optimization score using all samples. This statistic indicates how this resulting pipeline compares with the overall optimal pipeline in the complete dataset.

### Validation of somatic variants

In order to validate the robustness of the method we used both the optimized stringent (O) parameter values and a permissive (P) version of the algorithm (minimum number of forward and reverse reads supporting the variant=7 instead of 10). The permissive version allowed us to increase the number of the variants selected. We randomly selected for validation a subset of single nucleotide variants (O=154 out of 514, P=182 out of 758) and insertion-deletion mutations (O= 22 out of 227, P=16 out of 381) sequencing DNA amplicons containing the variants detected with our bioinformatic algorithm by targeted re-sequencing using AmpliSeq™ technology (Thermo Fisher Scientific, Waltham, MA, USA) according to the manufacturer’s specification. The AmpliSeq™ technology allows for a deep re-sequencing of the regions of interest, improving our ability to identify mutations correctly. We re-sequenced both tumor and control samples. Alternative alleles were validated if their frequency was ≥1%.

### Calculation of genetic divergence

We calculate genetic divergence between two samples as the number of mutations that are not shared between the two samples, divided by the total number of mutations in the union of the mutations detected in the two samples (expressed as a percentage). Divergence can only be reliably calculated if there are enough mutations to distinguish shared ancestry (mutations in common, sometimes called “public mutations”) from the evolution that has occurred after two populations last shared a common ancestor (private mutations). In order to reduce error in the divergence percentage calculation, we remove the samples with less than 5 total variants in the union of the SNVs called for both samples.

### Software availability

All software developed to carry out this study is distributed under the GPLv3 license. The implementation of the intratumor heterogeneity estimation pipeline—*ITHE*, can be found at https://github.com/adamallo/ITHE, scripts to carry out the cross-validation study and data analysis can be found at https://github.com/adamallo/ITHE_analyses, and the alternative implementation of our intratumor genetic heterogeneity pipeline using Mutect2 to call variants can be found at https://github.com/icwells/mutect2Parallel.

## Acknowledgments

This work was supported primarily by a CDMRP Breast Cancer Research Program Award BC132057, as well as NIH grants U54 CA217376, U2C CA233254, P01 CA91955, R01 CA170595, R01 CA185138 and R01 CA140657, and the Arizona Biomedical Research Commission grant ADHS18-198847. The findings, opinions and recommendations expressed here are those of the authors and not necessarily those of the universities where the research was performed or the funding bodies.

## Author Contributions

A.F. and D.M. developed the method and analyzed the data. S.M.R. contributed to the data analysis and implemented the alternative version of the bioinformatic pipeline that uses Mutect2. L.K. and T.H. collected patient’s clinical data and extracted the DNA from FFPE samples. A.H. review the histopathological status of the samples. A.F. and D.M. wrote the manuscript with support from C.C.M., J.R.M. and E.S.H. All authors discussed the results. C.C.M., J.R.M. and E.S.H supervised the project.

## Supplementary figures and tables

**Supplementary figure 1S:** Cross-validation.

**Supplementary figure 2S:** Sensitivity analysis.

**Supplementary figure 3S:** Genomic distribution of mutations. The cancer variants (O) were distributed across the entire exome.

**Supplementary table 1S:** Quantity of DNA sequenced (ng) and coverage.

**Supplementary table 2S:** Variants validation (O and P).

**Supplementary table 3S:** List of variants (O).

**Supplementary table 4S:** Gene panel, BRC83.

**Supplementary table 5S:** Mutect2 parameters.

